# Marker-free genome engineering in *Amycolatopsis* using the pSAM2 site-specific recombination system

**DOI:** 10.1101/2021.09.18.460463

**Authors:** Luísa D. F. Santos, Laëtitia Caraty-Philippe, Emmanuelle Darbon, Jean-Luc Pernodet

## Abstract

Actinobacteria belonging to the genus *Amycolatopsis* are important for antibiotic production and other valuable biotechnological applications such as biodegradation or bioconversion. Despite their industrial importance, tools and methods for the genetic manipulation of *Amycolatopsis* are less developed than in other actinobacteria such as *Streptomyces*. Moreover, most of the existing methods do not support convenient marker-free genome engineering. Here, we report the use of the pSAM2 site-specific recombination system for the efficient deletion of marker genes or large DNA regions in *Amycolatopsis*. For this purpose, we constructed a shuttle vector, replicating in *Escherichia coli* and *Amycolatopsis*, expressing the Xis and Int proteins from the *Streptomyces* integrative and conjugative element pSAM2. These proteins are sufficient for site-specific recombination between the attachment sites *attL* and *attR*. We also constructed two plasmids, replicative in *E. coli* but not in *Amycolatopsis*, for the integration of the recombination sites *attL* and *attR* on each side of a region targeted for deletion. We exemplified the use of these tools in *Amycolatopsis mediterranei* DSM 40773 by obtaining with high efficiency (>95%) a marker-free deletion of one single gene in the rifamycin biosynthetic gene cluster or of the entire 90-kb cluster.

**IMPORTANCE:** The genus *Amycolatopsis* is regarded as an important source of diverse specialized metabolites. Members of this genus are used in industry for the production of valuable antibiotics such as rifamycins or vancomycin. *Amycolatopsis* spp. also present a great interest for biotechnological applications such as biodegradation or bioconversion. Despite their importance, their genetic manipulation was somehow hampered by the lack of efficient tools. Here we report the successful use of the pSAM2 site-specific recombination system to construct unmarked deletion mutants, allowing marker recycling, or to create large deletions in *A. mediterranei* DSM 40773. The high efficiency of this site-specific recombination system and it possible application to other *Amycolatopsis* species open new opportunities for marker-free genome engineering in this genus.

## INTRODUCTION

Actinobacteria of the genus *Amycolatopsis* are the source of diverse bioactive specialized metabolites (1). Some species such as *Amycolatopsis mediterranei* and *Amycolatopsis orientalis* are for instance used industrially for the production of the medically important antibiotics rifamycins (2) and vancomycin (3, 4), respectively. *Amycolatopsis* spp. are also known for their biotechnological potential as agents of lignin degradation (5) and bioremediation (6) or for the fermentative production of aromatic compounds (7).

Despite the importance of the genus *Amycolatopsis*, the development of adapted genetic tools has lagged behind that for other actinobacterial genera such as *Streptomyces* (8, 9). For some time the introduction of DNA into *Amycolatopsis* has been a limiting factor, until this problem could be circumvented by the development of a method of direct transformation of *Amycolatopsis* mycelia (10) or by using intergeneric conjugation from *Escherichia coli* to introduce single-stranded DNA into the *Amycolatopsis* host (11, 12). Functional studies often rely on the inactivation/deletion of one or several genes. Recent attempts to use CRISPR-Cas9 based systems in *Amycolatopsis* reveal a toxicity effect of *Cas9* expression. A genome editing system based on other Cas protein was developed to obtain deletion mutants in *Amycolatopsis mediterranei* U32 (13), however this implicated the integration of the *Cas12a* gene and of the hygromycin resistance gene in the chromosome of *A. mediterranei* U32. A more common technique for gene inactivation is the replacement or deletion of the target gene by homologous recombination, the efficiency of which varies greatly depending on the strain (2), imposing the use of antibiotic resistance marker to select knockout mutants (14). As many *Amycolatopsis* species are naturally resistant to several antibiotics (15), the choice of antibiotic resistance marker might be limited and marker recycling is of particular interest when multiple genetic modifications have to be performed. To overcome this barrier some chromogenic screening systems associated to homologous recombination were developed and applied in *Amycolatopsis* (7) and in *Streptomyces* (16). These systems are based in the selection of simple recombinants, which integrated a suicide vector containing a reporter gene, using antibiotics resistance markers. The reporter gene confers a blue colour to the colonies of simple recombinants (in the case of *gusA* the presence of a specific substrate is necessary). Then, double recombinants are screened based in the loss of the blue colour, and white colonies are analysed to check if they contain the desired deletion or the initial genetic background.

Other techniques allowing marker free deletions and marker recycling based on site-specific recombination (SSR) have been developed to remove the antibiotic marker after knockout mutant selection in some actinobacteria (17). These techniques are based on the use of an antibiotic resistance gene cassette flanked by SSR sequences, which are recognized by cognate site-specific recombinases. The excision of the antibiotic cassette leaves a short scar (usually smaller than 50 bp) in the chromosome and can be designed to avoid polar effect that may occur if the antibiotic cassette is not removed (17). An example of well-characterized SSR system is that of pSAM2, an integrative element from *Streptomyces ambofaciens* (18). The integrase encoded by the *int* gene promotes the integration of pSAM2 into the chromosome by intermolecular SSR between the attachment (*att*) sites, *attP* carried by pSAM2 and *attB* site located in the bacterial chromosome. After integration, pSAM2 is flanked by the *attL* (left) and *attR* (right) sites. These sites can be themselves involved in intramolecular SSR leading to excision of pSAM2, an event requiring both the excisionase (encoded by the *xis* gene) and the integrase. Detailed studies have been carried out to precisely define the minimal sites required by the pSAM2 SSR system (19, 20). These findings were used for the construction of several excisable antibiotic resistance cassettes in which minimal *attL* and *attR* sites flank various antibiotic resistance genes (21). These selectable cassettes are used in *Streptomyces* to replace target genes by homologous recombination. After positive selection of the mutant strains, the resistance cassette can be efficiently excised by transiently expressing *xis* and *int* carried by an unstable vector. The resulting mutant strains are marker-free and contain a minimal *attB* site of 33, 34, or 35 bp. The size of the scar is dependent of the used excisable cassette, which is chosen in order to maintain the correct reading frame if the deletion is internal to the coding sequence(21). These cassettes can be used in *Amycolatopsis*, but without the possibility to excise them as the existing plasmids carrying the pSAM2 *int* and *xis* genes are unable to replicate in *Amycolatopsis*. The same is true for plasmids carrying other SSR systems that have be used in *Streptomyces*, as they all rely on replicons from pIJ101 or pSG5 (10).

Until now, few plasmids have been described in *Amycolatopsis* species (22), and only four of them (pMEA100, pMEA300, pA387 and pXL100) have been further studied. However, only the replicative vectors derived from the endogenous plasmids pA387 and pXL100 have been shown to replicate into several *Amycolatopsis* species (12, 23).

Here, we report the successful use of the pSAM2 SSR system to construct unmarked deletion mutants in *Amycolatopsis*, without leaving exogenous sequences within the genome, except for a short scar of 33 bp. For this purpose, we constructed a shuttle vector, replicating in *Escherichia coli* and *Amycolatopsis*, which is poorly maintained in *Amycolatopsis* in the absence of a selection pressure. This plasmid, derived from pRL60 (containing the short pA387 replicon) (24), carries the *xis* and *int* genes from pSAM2 and the *oriT* origin of transfer for interspecific conjugation. We also constructed two plasmids for the integration of the recombination sites *attL* and *attR* on each side of a region targeted for deletion. Then, with these newly constructed tools and also the previously constructed excisable antibiotic resistance cassettes (21) we demonstrated the efficiency of the pSAM2 SSR system to construct marker-free mutants and to generate large-scale deletions in *A. mediterranei* DSM 40773. These plasmids enrich the genetic toolbox available for genetic engineering of *Amycolatopsis* strains.

## RESULTS AND DISCUSSION

### Design of genetic tools for pSAM2 SSR system application

The application of the pSAM2 SSR system requires *cis-* and *trans*-acting elements and involves two steps (**Figure 1A**). First, the *cis* elements, the *attL* and *attR* recombination sites, are introduced in the genome via homologous recombination. Then, the *trans*-acting elements, the *xis* and *int* genes, are temporarily expressed to perform SSR and consequently the excision of the region flanked by *attL* and *attR*.

**Figure 1.**
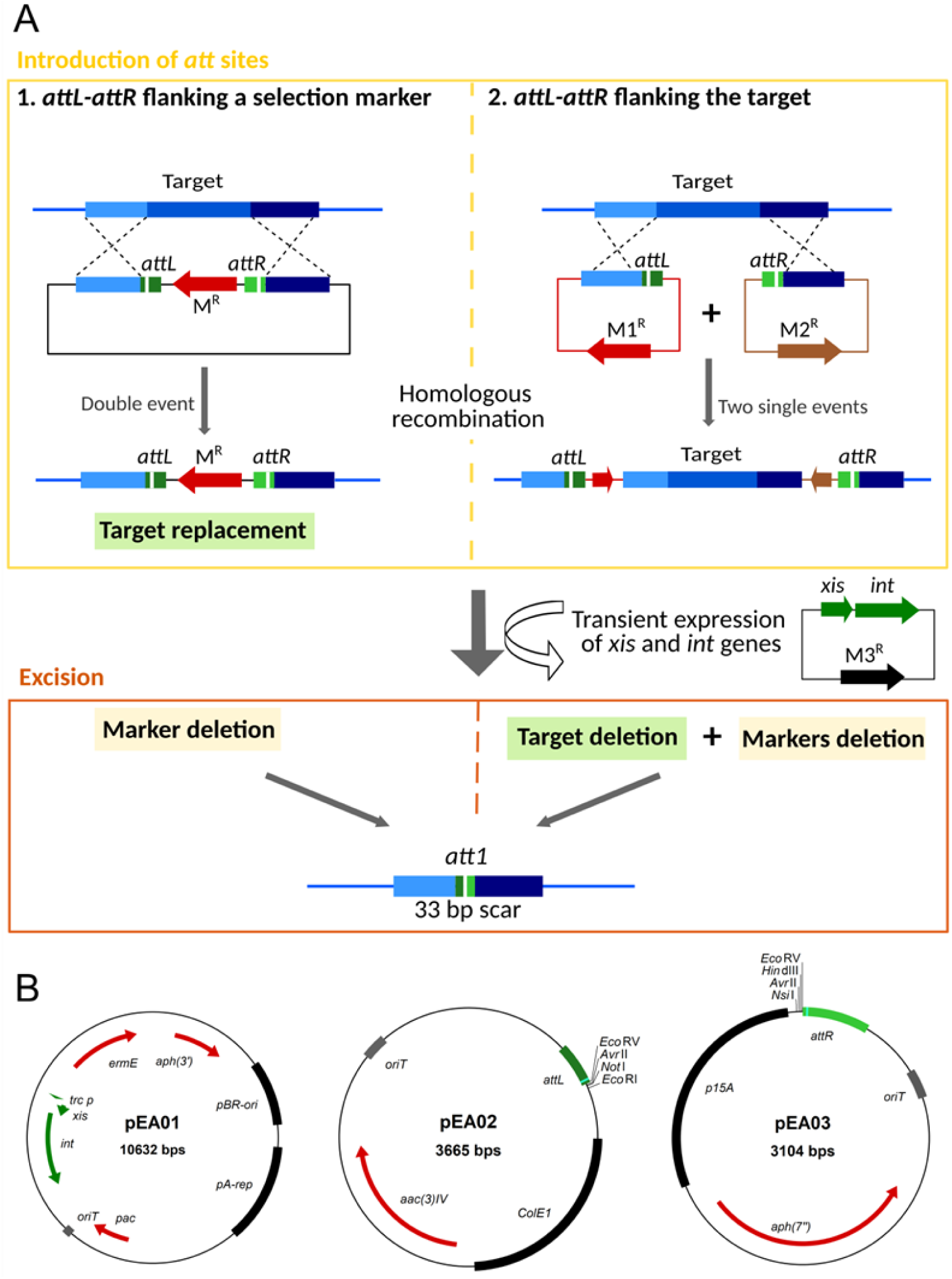
Principle of pSAM2 SSR system and genetic tools to create unmarked deletions. **A)** Examples of application of the pSAM2 SSR system (schematic representations not to scale). The SSR system requires *cis-*acting elements (*attL* and *attR* sites) and *trans-*acting elements (*xis* and *int* genes). First, the *att* sites are introduced in the genome via homologous recombination. The *attL* and *attR* sites are integrated in the genome by 1) a double event of homologous recombination in which the target is replaced by an excisable cassette/marker; or 2) two single events of homologous recombination after which the target is flanked by the *att* sites. Then the *xis* and *int* genes are introduced and temporarily expressed to perform the excision of the region flanked by *attL-attR* sites. **B**) Maps of the plasmids constructed in this study. All three plasmids are replicative in *E. coli*. pEA01 can replicate in *Amycolatopsis* while pEA02 and pEA03 are suicide vectors in *Amycolatopsis. aph(3’)*: kanamycin resistance gene; pBR-ori: pBR322 origin of replication; pA-rep: short replicon region of pA387; *pac*: puromycin resistance gene; *oriT*: origin of transfer; *int* and *xis*, integrase and excisionase gene of pSAM2; *trc*_*p*_: *trc* promoter for the expression of *xis* and *int* genes; *ermE:* erythromycin resistance gene; *attL*: left attachment site; ColE1: ColE1 origin of replication; *aac(3)IV:* apramycin resistance gene; *aph(7’’)*: hygromycin resistance gene; p15A: p15A origin of replication; *attR*: right attachment site.

The excision step requires only the transient expression of the *int* and *xis* genes. This can be achieved by cloning these genes in a vector self-replicating in *Amycolatopsis* which is rapidly lost in the absence of a selection pressure. For this purpose, we constructed a plasmid relying for replication in *Amycolatopsis* on the short form of the pA387 replicon contained in pRL60 (24), as this vector has been reported to be unstable in the absence of a selection pressure (25). The presence of the origin of transfer *oriT* allows the interspecific transfer of the plasmid from *E. coli* strain to *Amycolatopsis* species by conjugation, as wildly performed in actinobacteria. In our plasmid, three antibiotic resistance genes are present, conferring kanamycin, erythromycin or puromycin resistance, allowing its selection in various *Amycolatopsis* genetic backgrounds. The kanamycin marker is used for selection in *E. coli* strains. The *xis* and *int* genes are placed under the control of the promoter *trc*_p_, a promoter that efficiently expresses these genes in *Streptomyces* spp (21). The resulting plasmid possessing all these features was called pEA01 (**Figure 1B**). This plasmid is available from the Addgene repository (ID 170766).

The study of the vector replication, selection and stability was performed in *A. mediterranei* DSM 40773. After conjugative transfer of pEA01 to *A. mediterranei* DSM 40773, exconjugants resistant to erythromycin and puromycin were obtained. Of note, the kanamycin marker was not useful for this strain since *A. mediterranei* DSM 40773 is spontaneously resistant to kanamycin. The instability of pEA01 was assessed in absence of antibiotic selection: After 3 days of growth in media without antibiotic, 99.5% of clones (995 of 1000 analyzed) were sensitive to erythromycin indicating the loss of pEA01 vector. This confirmed that the short pA387 replicon from pRL60 is suitable to generate replicative but unstable vectors in *Amycolatopsis*, as required for transient expression of genes. The ability of pEA01 to replicate and its stability were also tested in two other *Amycolatopsis* strains, *Amycolatopsis sp. AA4* (26) and *Amycolatopsis sp*. ATCC39116 (5). The results obtained in these two strains are similar to those in *A. mediterranei* DSM 40773, *i*.*e*. pEA01 is able to replicate but unstable in the absence of selection pressure (**Table S1**).

Considering the introduction of *attL* and *attR* sites in the genome, two approaches can be used as described in **Figure 1A**. In the first one, the target gene(s) is replaced by an antibiotic resistance gene flanked by *attL* and *attR*. For this purpose, several excisable cassettes, previously constructed (21), are available for cloning in an *Amycolatopsis* suicide vector, in between regions identical to the upstream and downstream regions of the target gene. This approach was originally designed to inactivate a single gene by *in-frame* deletion for functional analysis (27). It can also be used to delete a few adjacent genes. Even if we used this approach to delete small (<30 kb) biosynthetic gene clusters (BGCs), such as the bicyclomycin BGC (28) and the congocindine BGC (29), the efficiency of the target replacement by homologous recombination decreases with the size of the target region. In this context, for large-scale deletions we propose a second approach in which *attL* and *attR* sites are successively introduced at the borders of the target region by homologous recombination. For this purpose, we designed two non-replicative vectors that share minimal sequence identity to avoid recombination between them. As both plasmids are conjugative, they nevertheless share an identical region corresponding to the *oriT* sequence (≈500 bp). This is the only long stretch of identity as we used different replicons (ColE1 and p15A) for replication in *E. coli* and different antibiotic resistance genes (apramycin and hygromycin resistance). The two vectors pEA02 and pEA03, contain the *attL* and *attR* sites, respectively (**Figure 1B**). They both harbor a multiple cloning site for the easy cloning of the sequences required for homologous recombination between the vector and the host genome. These vectors are suitable for introduction of *attL* and *attR* sites at any target borders by a single event of homologous recombination. Detailed instructions to clone homologous regions into these vectors are provided in **Figure S1**. pEA02 and pEA03 are available from Addgene (ID 172193 and ID 172194, respectively).

The set of three vectors and the efficiency of each approach were thereafter evaluated for single gene or large-scale deletions in *Amycolatopsis*. We used as target the rifamycin biosynthetic gene cluster of *A. mediterranei* DSM 40773.

### Cassette excision

To demonstrate the application of the pSAM2 SSR system to create unmarked mutants by cassette excision, we first targeted the *rifK* gene, encoding a AHBA (3-amino-5-hydroxy benzoic acid) synthase (2). AHBA is the starter unit of rifamycin biosynthesis, so the inactivation of *rifK* abolishes the production of the precursor and consequently of rifamycin. The phenotype of the mutant strains could be easily tested by bioassays.

The strategy employed for the obtention of marker-free *rifK* mutants is summarized in **Figure 2A**. First *rifK* was replaced by an excisable hygromycin resistance cassette (*att1Ωhyg*) following double homologous recombination events between the non-replicative plasmid pRIF05 and the host genome. For that, pRIF05 containing the excisable cassette between the upstream and the downstream homologous regions was introduced into the wild-type strain by conjugation. Hygromycin resistant exconjugants were selected and then tested for sensitivity/resistance to apramycin. Of the 300 hygromycin resistant clones tested only 20 were sensitive to apramycin, indicating that double homologous recombination events are rather rare (less than 10%). These hygromycin-resistant and apramycin-sensitive clones were then checked by PCR on genomic DNA. The verification is based on the amplification of the junctions formed between the upstream or the downstream region of *rifK* and the *att1Ωhyg* cassette. As expected only the clone of interest allowed the amplification of DNA fragments with the expected size (PCR 1 and PCR 2 in **Figure 2B**). The results are shown for one clone, representative of 20 clones analyzed. The resulting clones were called *A. mediterranei* DSM 40773 *ΔrifK::att1Ωhyg*, and three of them were used in further steps and for phenotypic analysis.

**Figure 2.**
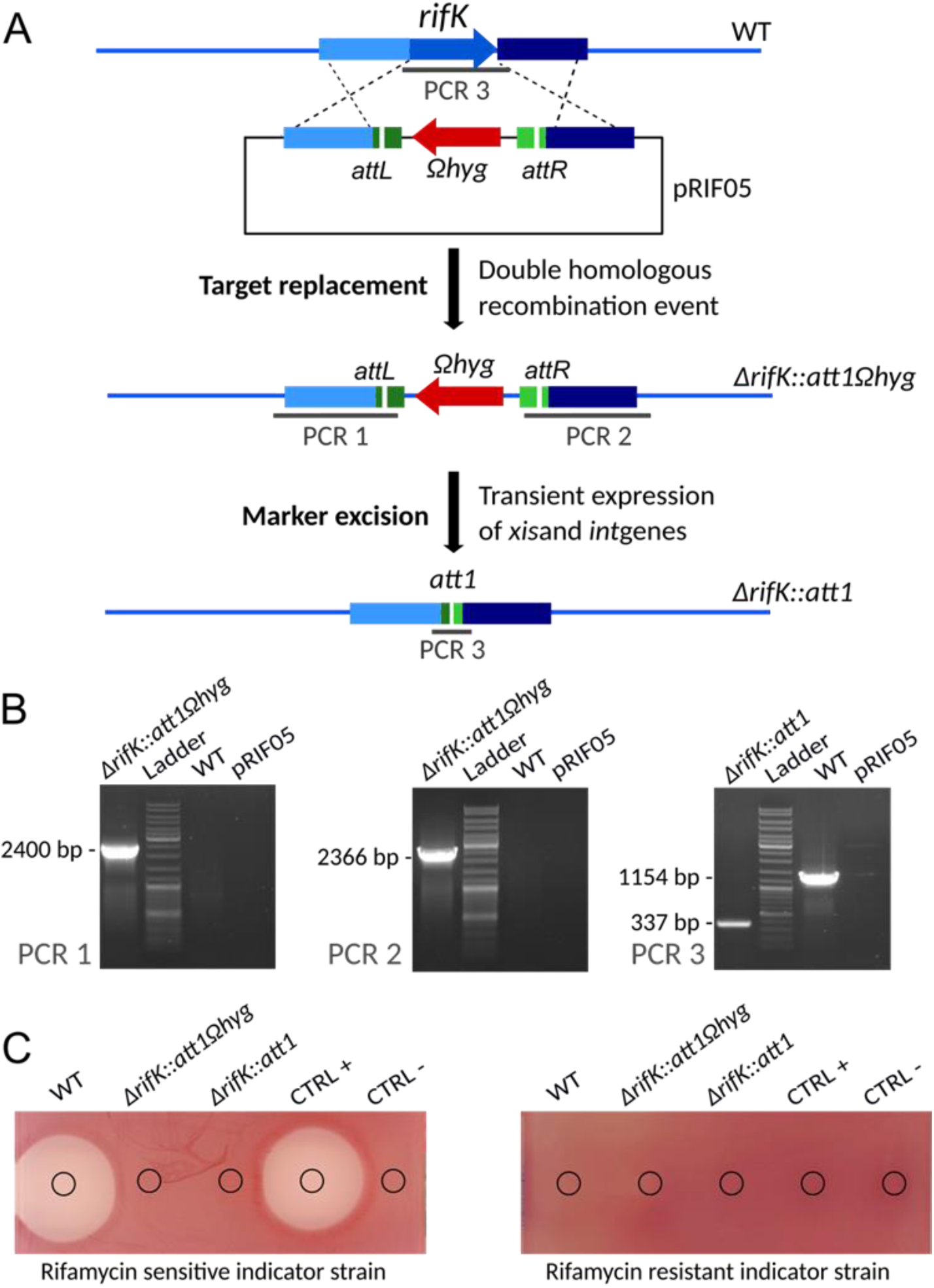
Construction of an unmarked *rifK* deletion mutant using an excisable cassette. **A)** Schematic representation (not to scale) of the successive steps. First the *rifK* gene is replaced by the *att1Ωhyg* cassette via a double homologous recombination event. Then the *att1Ωhyg* cassette is excised following the introduction of pEA01, resulting in an unmarked deletion mutant containing a 33-bp scar (*att1*). **B)** PCR verification of the strains at different stages of the construction. Verifications are based on three PCRs. The extent of the expected amplicons for each of the PCR is indicated in panel A. The amplicons sizes are indicated on the left side of the pictures. Lanes: *ΔrifK::att1Ωhyg* – genomic DNA from one of the *A. mediterranei* DSM43770 *ΔrifK::att1Ωhyg* clones, representative of all analyzed clones; Ladder – GeneRuler DNA Ladder (SM0331 Thermo Scientific); *ΔrifK::att1* – genomic DNA from one of the *A. mediterranei* DSM43770 *ΔrifK::att1* clones, representative of all analyzed clones WT – genomic DNA from *A. mediterranei* DSM43770 wild-type strain; pRIF05 – Plasmid pRIF05. **C)** Bioassays. The antibacterial activity of the culture supernatants from the wild-type and the mutant strains of *A. mediterranei* DSM43770 was assayed against *S. aureus* HG003 (sensitive to rifamycin) and its *rpoB*_H481Y_ mutant derivative (resistant to rifamycin). MP5 with or without 5 µg/mL of rifamycin SV was used as positive or negative control, respectively.

In the second step, the excision of the *att1Ωhyg* cassette was performed by introducing the plasmid pEA01 in three independent *ΔrifK::att1Ωhyg* clones. The erythromycin resistant exconjugants obtained were then screened for resistance/sensitivity to hygromycin to check the loss of the hygromycin cassette. Over the 150 (50 from each parental clone) clones tested 147 were sensitive to hygromycin indicating that the excision occurs with a frequency of 98%. Eighteen of these hygromycin sensitive clones (6 from each parental clone) were genetically verified by PCR, targeting the scar region formed by the excision of *Ωhyg* cassette. As expected, a smaller fragment was amplified with genomic DNA of the sensitive hygromycin clones when compared to the 1154 bp fragment obtained with the wild-type genomic DNA. The results obtained for PCR 3 are also shown in **Figure 2B** for one clone, representative of the 18 clones tested. The sequence analysis of the 337 bp amplicon, confirmed the excision of the *Ωhyg* cassette and the formation of a 33 bp scar, corresponding to the *att1* sequence (21). Moreover, these clones became sensitive to erythromycin, indicating that pEA01 has been lost during cultivation in the absence of erythromycin. The resulting clones were called *A. mediterranei* DSM 40773 *ΔrifK::att1*, and three of them (one from each parental clone) were used for phenotypic analyses.

Phenotypic analysis, presented in **Figure 2C**, was performed using a bacterial growth inhibition assay. The results shown that *ΔrifK::att1Ωhyg* and *ΔrifK::att1* strains did not produce rifamycin contrarily to the wild-type strain, confirming that this gene is essential for antibiotic production.

So far, we validated the application of pSAM2 SSR system for cassette excision to generate unmarked *ΔrifK* mutants in *A. mediterranei* DSM 40773. Moreover, we also used this approach to successfully obtain deletion mutants in *Amycolatopsis sp*. AA4 and *Amycolatopsis sp*. ATCC 39116 (See Table S1 for excision efficiency).

### Region excision

The complete rifamycin gene cluster was targeted to demonstrate the use of the pSAM2 SSR system to generate marker-free large-scale deletions. The rifamycin biosynthetic gene cluster comprises 42 genes covering a region of about 90 kb (30). The strategy employed to delete this cluster is shown in **Figure 3A**. First the *attL* and *attR* sequences were integrated at the cluster’ extremities via homologous recombination. In a second step, the excision of the region flanked by *attL-attR* sites and encompassing the rifamycin gene cluster was performed using the pEA01 vector.

**Figure 3.**
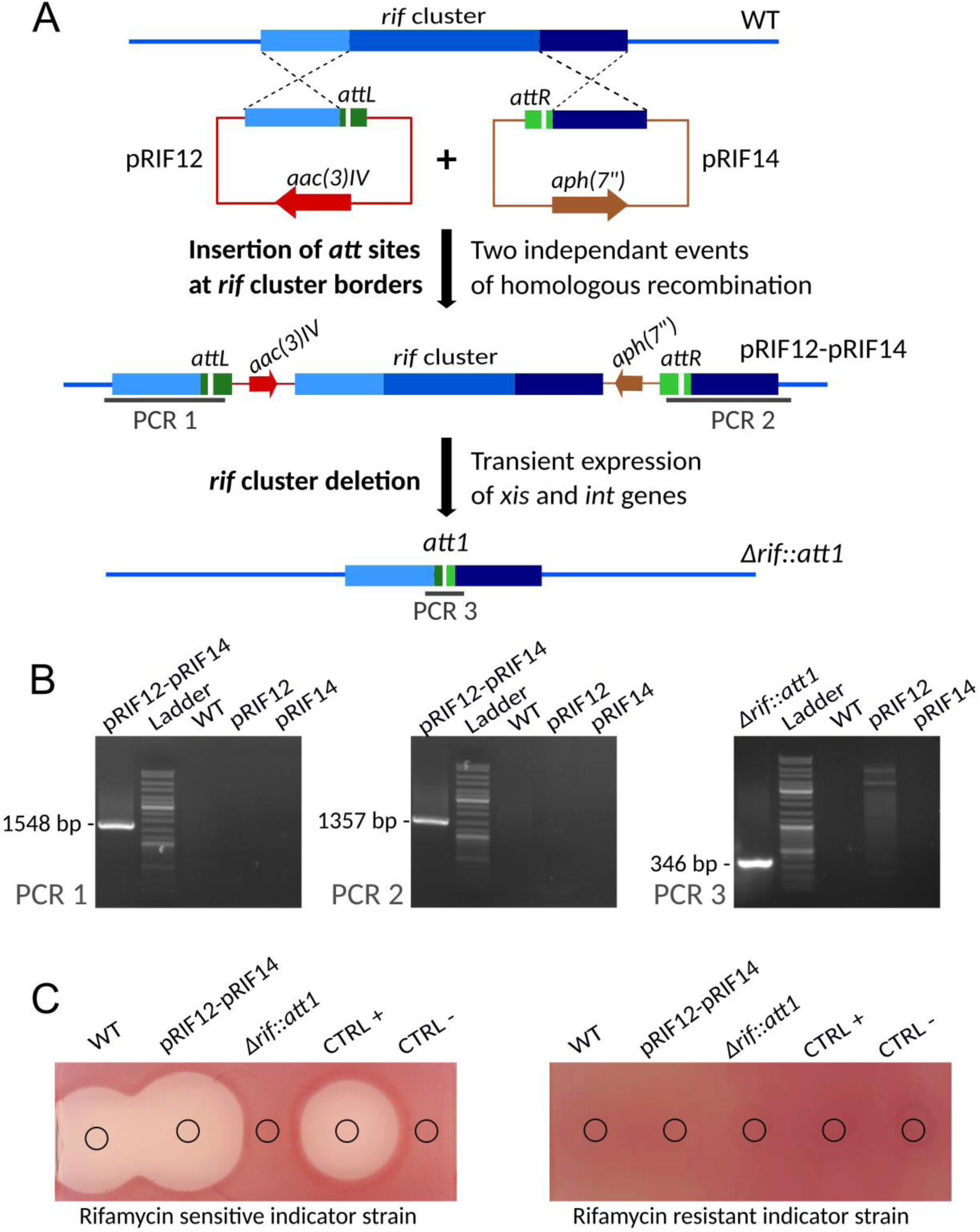
Construction of an unmarked *rif* cluster deletion mutant. **A)** Schematic representation (not to scale) of the successive steps. First the *attL* and *attR* sequences are successively integrated upstream and downstream of the *rif* cluster via two single homologous recombination events. Then the complete region between the *attL* and *attR* sites is excised following the introduction of pEA01, resulting in an unmarked *rif* cluster deletion mutant containing a 33-bp scar (*att1*). **B)** PCR verification of the strains at different stages of the construction. Verifications are based on three PCRs. The extent of the expected amplicons for each of the PCR is indicated in panel A. Lanes: pRIF12-pRIF14 – genomic DNA of *A. mediterranei* DSM43770 harboring pRIF12 and pRIF14, representative of all analyzed exconjugants; Ladder – GeneRuler DNA Ladder (SM0331 Thermo Scientific); WT – genomic DNA of *A. mediterranei* DSM43770 wild-type strain; pRIF12 – Plasmid pRIF12; pRIF14 – Plasmid pRIF14; *Δrif::att1* – genomic DNA of *A. mediterranei* with a deletion of the *rif* cluster. **C)** Bioassays. The antibacterial activity of culture supernatants from the wild-type strain *A. mediterranei* DSM43770, the strain harbouring pRIF12 and pRIF14, and the mutant strain *Δrif::att1* was assayed against *S. aureus* HG003 (sensitive to rifamycin) and its *rpoB*_H481Y_ derivative (resistant to rifamycin). MP5 with or without 5 µg/mL of rifamycin SV was used as positive or negative control, respectively.

For this purpose, fragment identical to the region upstream and downstream of the rifamycin cluster were cloned into pEA02 and pEA03, yielding pRIF12 and pRIF14, respectively. These plasmids were then transferred into the wild-type strain by successive conjugations. After conjugative transfer of pRIF14 into the wild-type strain, clones resistant to hygromycin were selected. These clones were then verified by PCR on genomic DNA to confirm the integration of pRIF14 at the downstream extremity of rifamycin cluster. In a second step, pRIF12 was transferred into the strain already carrying the pRIF14, and clones resistant to apramycin and hygromycin were selected. Then, as previously described for strains carrying the pRIF14, these clones were verified by PCR. The results obtained confirmed the integration of pRIF12 and pRIF14 at the expected chromosomal sites (upstream and downstream to the rifamycin gene cluster) as presented in **Figure 3B** for PCR 1 and PCR 2. The resulting clones were called *A. mediterranei* DSM 40773-pRIF12-pRIF14 and were used in further steps and for phenotypic analysis. From the literature, it appears that the efficiency of homologous recombination in *Amycolatopsis* spp is strain dependent (2, 13). Thus it might be preferable to use rather large recombination regions to integrate the *attL* and *attR* sequences.

The excision of the rifamycin gene cluster was performed in two independent *A. mediterranei* DSM 40773-pRIF12-pRIF14 clones, using the plasmid pEA01. The erythromycin resistant clones obtained after conjugation of pEA01 were then screened for resistance/sensitivity to hygromycin and/or apramycin to check the loss of hygromycin and/or apramycin cassette. Over the 100 tested exconjugants (50 from each parental clone) that received pEA01 vector, 95 were sensitive to both antibiotics indicating that the excision occurs with a frequency of 95%. Twelve of these sensitive clones (6 from each parental clone) were verified by PCR, targeting the scar region formed after excision (PCR 3 in **Figure 3B**). As expected, a DNA fragment of 346 bp, containing the *att1* sequence (21), was amplified for all the 12 clones tested. Moreover, these clones became sensitive to erythromycin, indicating that pEA01 has been lost during cultivation in the absence of erythromycin. The resulting clones were called *A. mediterranei* DSM 40773 *Δrif::att1*, and three of them were used for phenotypic analyses.

As previously described for *rifK* mutants, bacterial growth inhibition assay results confirmed that the *Δrif::att1* strains did not produce rifamycin, as shown in **Figure 3C**. The integration of pRIF12 and pRIF14 at the border of the cluster does not affect the production of rifamycin.

This experiment validates the functionality of the set of vectors developed in this work and the application of the pSAM2 SSR system to generate large-scale deletions in *A. mediterranei* DSM 40773.

### Concluding remarks

The genetic tools that we constructed allow the application of the pSAM2 SSR system in *A. mediterranei* to generate marker-free small or large deletions. The SSR between *attL* and *attR* leading to the excision of the region flanked by these sites is highly efficient (> 95%). The efficiency is roughly the same whether these sites are close (98% efficiency with excisable cassettes) or quite distant (*e*.*g*. 95% efficiency for sites separated by 90 kb). These tools allow the use of resistance marker for steps where a selection is helpful and the subsequent removal of these markers by SSR to obtain marker-less deletion mutant strains. This allows the recycling of resistance markers for successive rounds of genetic engineering, an advantage for a genus for which the number of usable resistance markers is limited. Moreover, the absence of heterologous antibiotic resistance genes might be important for biotechnological applications. In addition, unmarked *in-frame* gene deletion prevents polar effects on the expression of downstream genes.

The application of pSAM2 SSR system to generate unmarked deletion mutants requires the introduction of several vectors into *Amycolatopsis*: 1) the introduction of either a suicide vector for gene replacement by a selection marker or two suicide vectors for the introduction of *attL* and *attR* at the borders of the target region; 2) the introduction of an unstable self-replicative vector for excision. Thus, this process might seem more time-consuming than the ones using double homologous recombination techniques associated to a chromogenic screen (7, 16), which require the introduction of a single vector. In these systems, the integration of a suicide vector by simple crossing over is selected using a resistance marker and then the loss of the vector, after a second event of homologous recombination, can be screen by the colour of the colonies. However, obtaining clones in which a second event of homologous recombination had occurred may require several rounds of sporulation in non-selective media (7). These clones correspond either to a reversion to wild-type or to the desired deletion. The main advantage of the approach described in this work is that the steps based on homologous recombination are always associated with the ability to select the expected event by antibiotic resistance. The only step for which there is no selection but only screening is the excision step. Using the pSAM2 SSR system allows to obtain a very high frequency (> 95%) of excision, so that a selection for the desired mutant is not required. Therefore, the genetic tools developed here enriches the toolbox available for marker-free genome engineering of *Amycolatopsis* strains and might facilitate the exploration and exploitation of their biotechnological potentialities. Depending on the efficiency of the homologous recombination in the *Amycolatopsis* strain used and on the size of the deletion to be created, it might preferable to use the chromogenic systems previously described (7, 16) or the system described here. It should also be noted that the plasmids pEA02 and pEA03 might be used in *Streptomyces* spp. in combination with the *Streptomyces* replicative vectors expressing Xis and Int (pOSV236 (31), pOSV507 or pOSV508 (21)) to obtain large-scale deletions.

## MATERIALS AND METHODS

### Bacterial strains, cultivation conditions and strain manipulation

All strains used in this study are listed in **Table 1**. *E. coli* strains were grown at 37°C in LB. When required, antibiotics were added to *E. coli* cultures in liquid (or on solid) medium at the following concentrations: ampicillin, 50 µg/ml (or 100 µg/ml); apramycin, 25 µg/ml (or 50 µg/ml); hygromycin, 50 µg/ml (or 150 µg/ml); kanamycin, 25 µg/ml. *A. mediterranei* strains were grown at 30°C on GYM agar medium (32) for sporulation before the preparation of spore stocks, in TSB (Tryptic Soy Broth, Becton Dickinson) for DNA extraction and in MP5 for rifamycin production. Conjugations between *E. coli* ET12567 harboring pUZ8002 (or pUZ8003) and *A. mediterranei* were carried out according to Kieser *et al* (34) using MS medium complemented with 10 mM CaCl_2_ (35), instead of MgCl_2_. An overlay (3 ml) of soft nutrient agar (Nutrient Broth, Oxoid, with 0.8% agar) containing 25 µg/ml nalidixic acid and the appropriate antibiotics for the selection of exconjugants was added after overnight incubation of the conjugation plates. The plates were incubated for 5–7 days until exconjugants clones became visible. Antibiotic concentration used for *A. mediterranei* were as follows: apramycin, 50 µg/ml; erythromycin, 75 µg/ml; hygromycin, 75 µg/ml.

**Table 1.**
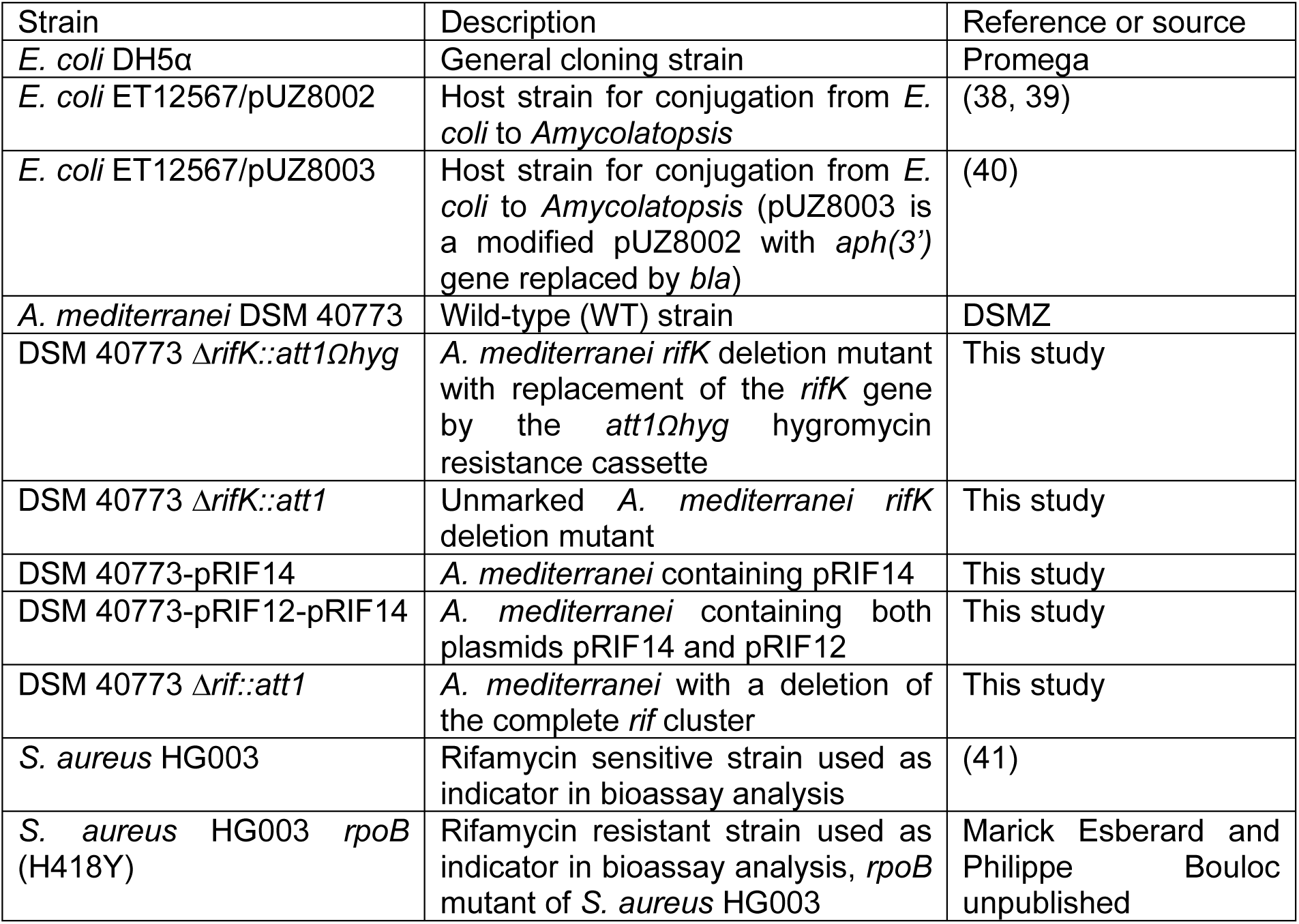
Strains used in this study.

### Plasmids and DNA manipulations

All plasmids used in this study are listed in **Table 2**. Details about the plasmids constructed for this study are given below. Amplification of DNA fragments for cloning was carried out using the high-fidelity DNA polymerase Q5 (New England BioLabs: NEB). Taq polymerase (Qiagen) was used for verification PCRs. All oligonucleotides used in this study were provided by IDT and are listed in **Table S2**. Restriction and modification (ligase, kinase, etc.) enzymes were purchased from NEB or ThermoScientific. Plasmid DNA extraction from *E. coli* was performed using NucleoSpin Plasmid kit from Macherey-Nagel. DNA fragments were purified from agarose gels using the NucleoSpin Gel and PCR Clean-up kit from Macherey-Nagel. Genomic DNA extractions from *Amycolatopsis* and *E. coli* transformations were performed according to standard procedures (34, 36).

**Table 2.**
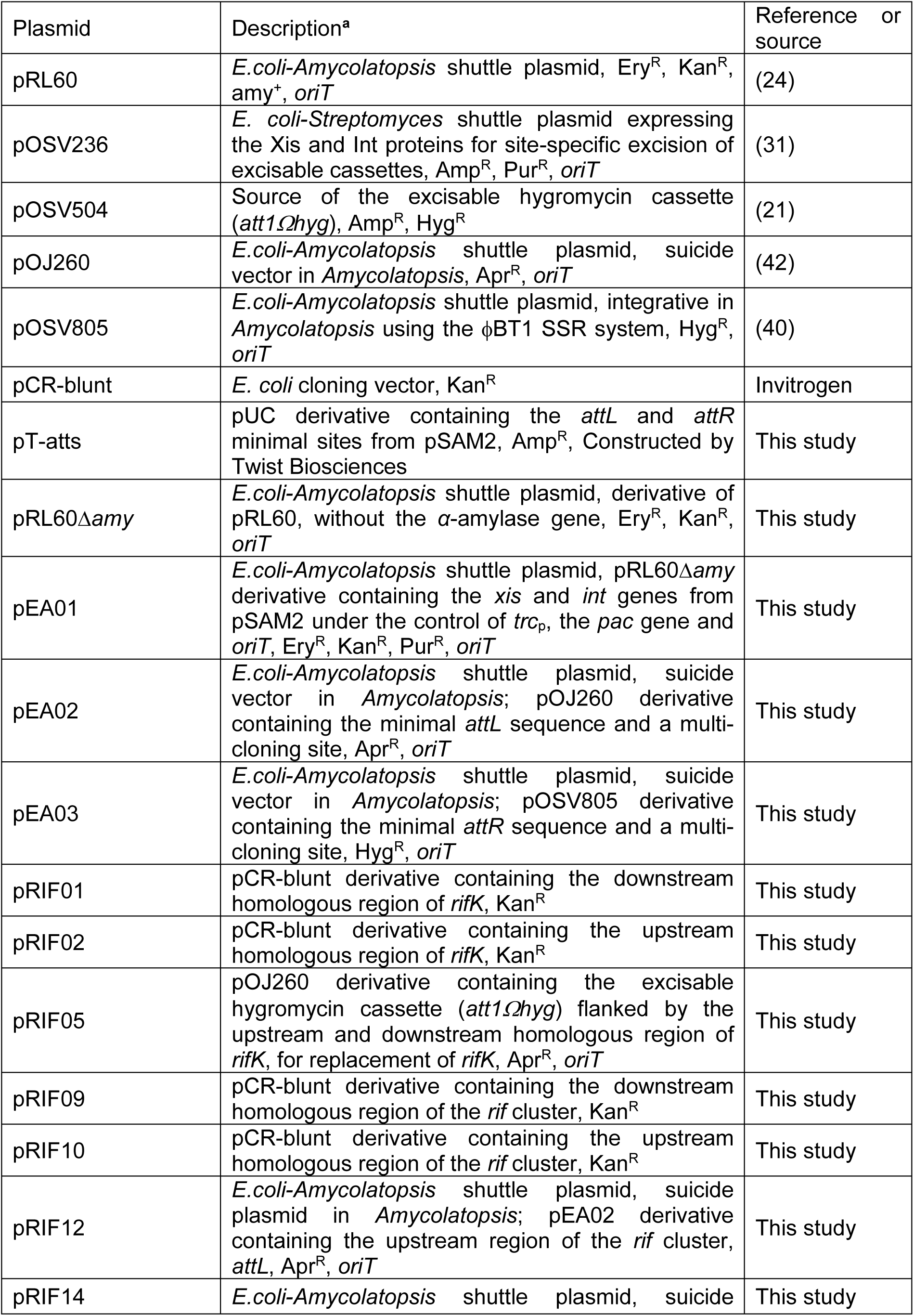
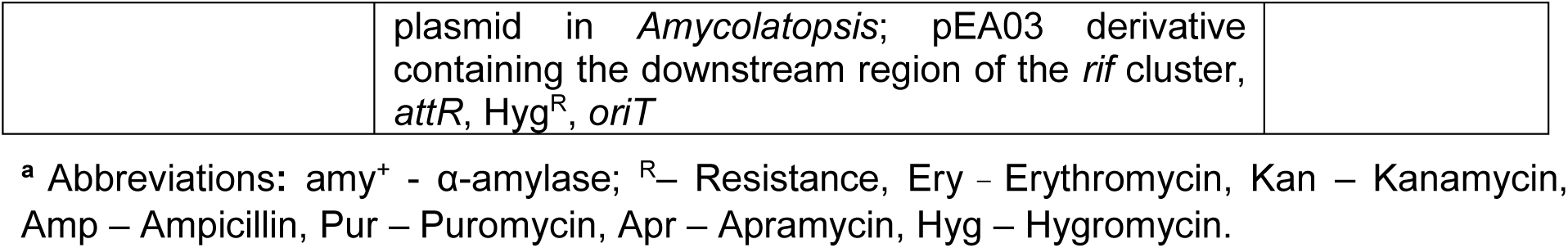
Plasmids used in this study.

### Plasmid construction

#### (i) Construction of the plasmid pEA01 expressing Xis and Int

pRL60 was digested by *Eco*RI and DNA fragments of 6 kb (*aph(3’)* gene, pBR*-*origin and pA-rep) and 1,8 kb (*ermE* gene) were then purified and ligated, yielding pRL60Δ*amy*. The DNA fragment containing the *xis* and *int* genes under control of *trc*_p_, the puromycin resistance gene *pac* and the origin of transfert *oriT* was isolated from pOSV236 by digestion with *Not*I and *Ssp*I. This 3 kb fragment was then blunt-ended using Klenow enzyme and cloned into pRL60Δ*amy* linearized by *Eco*RV and dephosphorylated, yielding pEA01. The construction was verified by restriction enzyme digestion and sequencing.

#### (ii) Construction of the plasmids pEA02 and pEA03 carrying *attL* and *attR*

A DNA fragment containing the *attL* and *attR* sites surrounded by desired restriction sites was chemically synthesized and cloned into a high copy number plasmid by Twist Bioscience, yielding pT-atts. A *Bam*HI/*Eco*RI fragment containing the *attL* sequence flanked by the *Eco*RV, *Avr*II and *Not*I restriction sites was isolated from pT-atts and cloned into pOJ260 linearized by *Bam*HI and *Eco*RI, yielding pEA02. A *Pst*I*/Nsi*I fragment containing the *attR* sequence flanked by the *Eco*RV, *Hin*dIII and *Avr*II restriction sites was isolated from pT-atts and ligated to a 2.8 kb fragment obtained by digestion of pOSV805 by *Pst*I and *Nsi*I, yielding pEA03. It should be noted that the *int* gene from ϕBT1, present in pOSV805, is no longer present in pEA03. Thus pEA03 is a suicide vector in *Amycolatopsis*, as pEA02. Both plasmids were verified by restriction analyzes and partial sequencing.

#### (iii) Construction of pRIF05 for the inactivation of *rifK*

The regions upstream and downstream (≈2 kb each) of *rifK* were amplified by PCR using the following couple of primers: LS151/LS152 and LS153/LS154, respectively. Each PCR product was cloned into pCR-blunt, yielding pRIF02 and pRIF01. Effective cloning of the fragments in pCR-blunt was controlled by restriction analysis and the integrity of the insert was verified by sequencing. The downstream region was then isolated from pRIF01 as a *Hin*dIII/*Pvu*II fragment, while the upstream region was isolated from pRIF02 as a *Bam*HI/*Pvu*II fragment. The *att1Ωhyg* cassette was isolated from pOSV504 by *Eco*RV. The three DNA fragments were then cloned together into pOJ260 digested by *Bam*HI and *Hin*dIII, yielding pRIF05.

#### (iv) Construction of pRIF12 and pRIF14 for deletion of the rifamycin gene cluster

The regions upstream (1.2 kb) and downstream (1 kb) of the rifamycin (*rif*) biosynthetic gene cluster were amplified using the pairs of primers: LS155/LS156 and LS157/LS158 respectively. Each PCR product was cloned in pCR-blunt, yielding pRIF10 and pRIF09. Effective cloning of the fragments in pCR-blunt was controlled by restriction analysis and the integrity of the insert was verified by sequencing. The *Eco*RV/*Not*I fragment of pRIF10 containing the upstream region of the *rif* cluster was then cloned into pEA02 digested by the same enzymes yielding pRIF12. In the same way, to construct pRIF14, the downstream region of the *rif* cluster was isolated from pRIF09 as a *Nsi*I/*Eco*RV fragment and was then cloned into pEA03 digested by the same enzymes.

### Construction and verification of *A. mediterranei* DSM 40773 mutant strains

#### (i) Marker-free *rifK* mutants

Firstly, pRIF05 was transferred to *A. mediterranei* wild-type by conjugation. Hygromycin resistant exconjugants were selected and then screened for apramycin sensitivity to obtain clones in which the *rifK* gene was replaced by the *att1Ωhyg* cassette. The genetic organization of hygromycin-resistant and apramycin-sensitive clones was checked by PCRs targeting the upstream and the downstream junctions formed by the replacement of the *rifK* gene by the *att1Ωhyg* cassette. These PCRs were performed using the following couples of primers: LS182/LS161 and LS188/LS189, respectively. For excision of the *att1Ωhyg* cassette, the plasmid pEA01 was introduced into three independent *ΔrifK::att1Ωhyg* clones. Erythromycin resistant exconjugants were selected. Hygromycin sensitive clones were then screened and the loss of plasmid pEA01 was finally obtained by successive subculturing on medium without erythromycin. The excision of the cassette and the deletion of the *rifK* gene was checked by PCR using primers hybridizing upstream and downstream of the *rifK* gene (LS172 and LS173 respectively). For each mutant clone, the presence of the scar was checked by sequencing the PCR product.

#### (ii) Large-scale deletions

Firstly, pRIF14 was transferred to *A. mediterranei* wild-type by conjugation. Hygromycin resistant exconjugants were selected. The integration of the plasmid pRIF14 was checked by PCR using primers LS190 and LS71 targeting the downstream junction formed by the integration of pRIF14 within the region downstream of the *rif* cluster. pRIF12 was then introduced into the strain already harboring pRIF14. Apramycin and hygromycin resistant exconjugants were selected. The integration of the plasmid pRIF12 was checked by PCR using primers LS191 and LS70 targeting the upstream junction formed by the integration of pRIF12 within the region upstream of the *rif* cluster. Finally, the excision of the complete cluster was performed as previously described for the excision of the hygromycin cassette. The deletion of the complete *rif* cluster was checked by PCR using primers hybridizing upstream and downstream of the *rif* cluster (LS220 and LS221 respectively) and allowing the amplification of the scar region formed after the excision. For each mutant clone, the presence of the scar was checked by sequencing the PCR product.

### Rifamycin production

MP5 liquid medium (75 ml of medium in 500 ml baffled Erlenmeyer flasks) was inoculated with 5×10^7^ spores of the mutant or the wild-type strains. Cultures were incubated for 10 days at 30°C under agitation (180 rpm). Cultures were centrifugated and the supernatants used for the assay of the antibacterial activity.

### Antibacterial activity assays

Antibacterial activity assays were performed using two *Staphylococcus aureus* strains as indicator (one strain sensitive to rifamycin and the other resistant). Each *S. aureus* strain was grown overnight in LB at 37°C and used to inoculate molten Antibiotic medium 5 (Becton Dickinson). Each indicator plate was loaded with 100 µl of *A. mediterranei* supernatants and then incubated overnight at 37°C for growth inhibition analysis. The growth inhibition of strains by rifamycin was verified with 100 µl of MP5 containing or not rifamycin SV (5 mg/ml). The contrast between regions of normal and inhibited growth was enhanced using tetrazolium red (Sigma) as described by Pattee (37). These assays were performed at least three times for each mutant strain.

## SUPPLEMENTAL MATERIAL

Supplemental material for this article is found after the “references” section.

## ACKNOWLEDGMENTS

We thank Prof. Rup Lal for the kind gift of the plasmid pRL60 and Marick Esberard and Dr. Philippe Bouloc for the kind gift of *S. aureus* strains. We thank Jorddy Benoist, Anaïs da Costa and Gaëtan Pavard for their technical help and motivation during their internships. We thank Prof. Stéphanie Bury-Moné for critical reading of the manuscript and helpful discussions. This study was funded by the ANRT (CIFRE-2018/0227). The funders had no role in the design of the study and collection, analysis, and interpretation of data and in writing the manuscript.

## SUPPLEMENTAL MATERIAL

**Figure S1.**
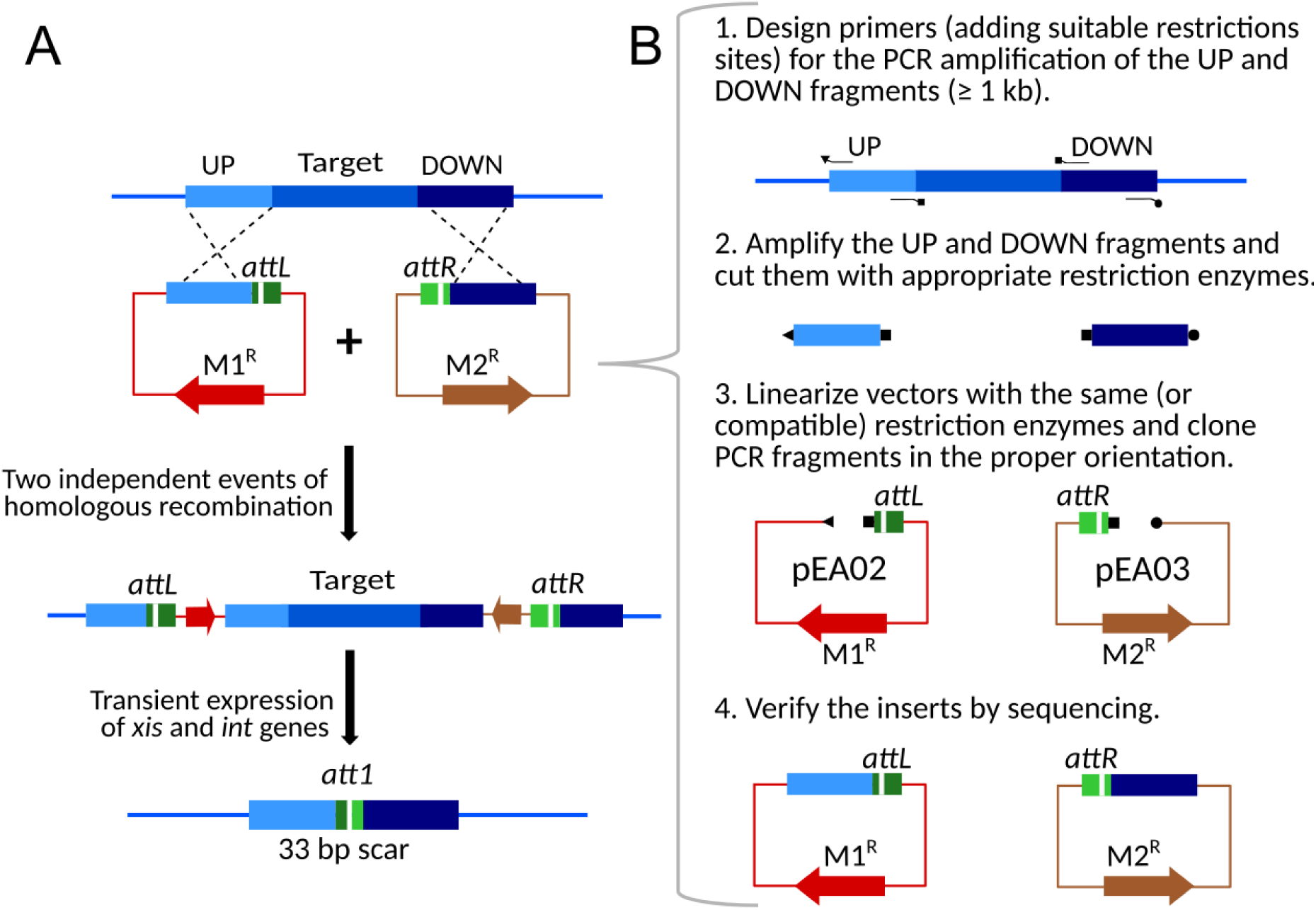
Large-scale marker-free deletions using the pSAM2 SSR system. **A)** Schematic representation of the successive steps for large region excision (not to scale). First, the minimal *attL* and *attR* sites are successively introduced at the borders of the target region by homologous recombination. *att* sites are represented by a green rectangle in which the white line represents the crossover sequence (TCGGG) where SSR takes place. For proper excision of the large region (target region, vectors and markers used to introduce the *att* sites), these *att* sites should be external to this large region. Then, the deletion is performed by the expression of the *xis* and *int* genes. The intramolecular SSR between minimal *attL* and *attR* sites leads to the formation of the *att1* site. **B)** Main steps to clone the homologous fragments (identical fragments to the upstream (UP) and the downstream (DOWN) regions) into pEA02 and pEA03 for the correct integration of *att* sites at the target borders. ◂ Represents *Avr*II, *Not*I, *Eco*RI (or compatible) restriction sites; ◼ Represents *Eco*RV (or any blunt end) restriction site or no restriction site (for blunt end PCR products) ● Represents *Hin*dIII, *Avr*II, *Nsi*I (or compatible) restriction sites.

**Table S1.**
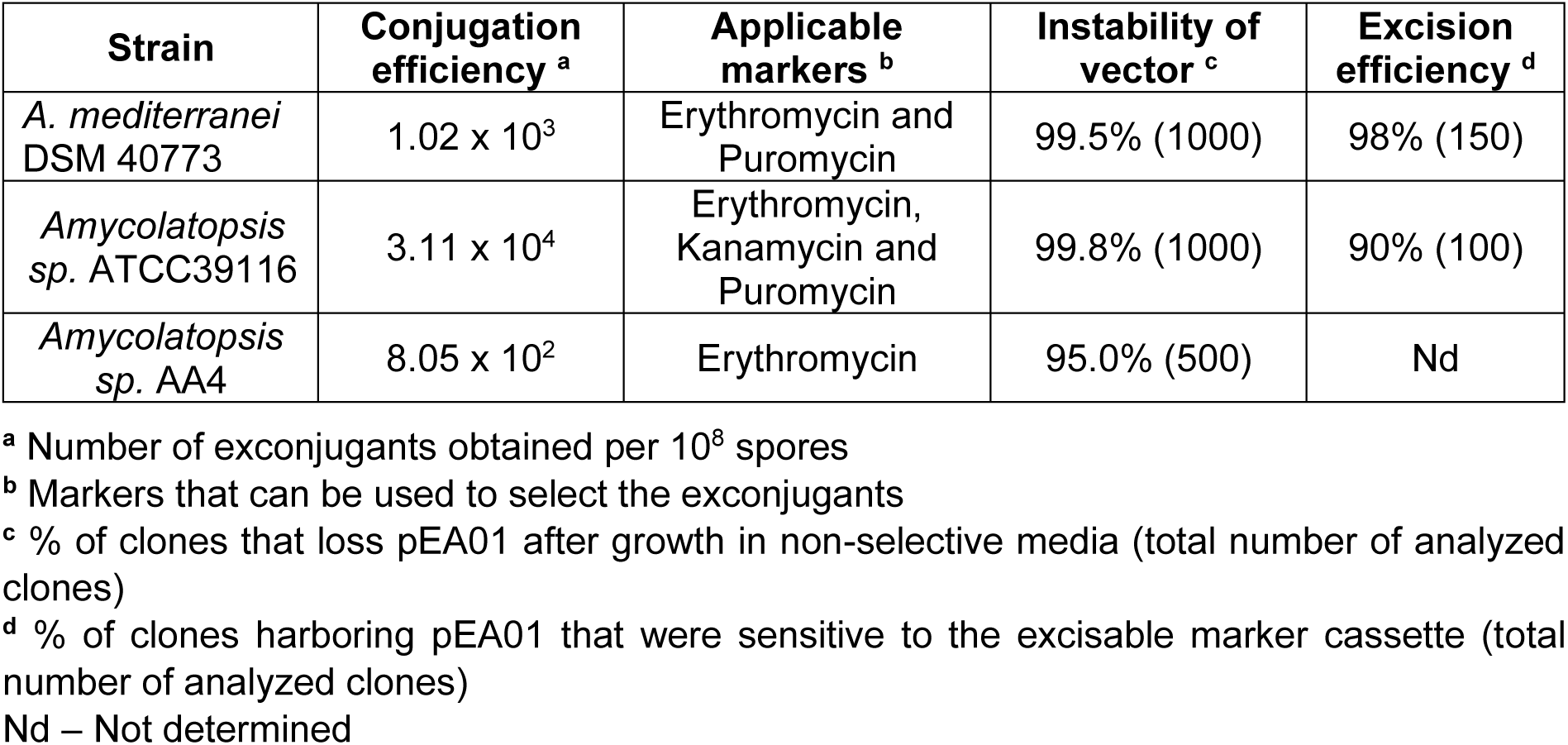
Analyses of the pEA01 functionality in *Amycolatopsis* species.

**Table S2.**
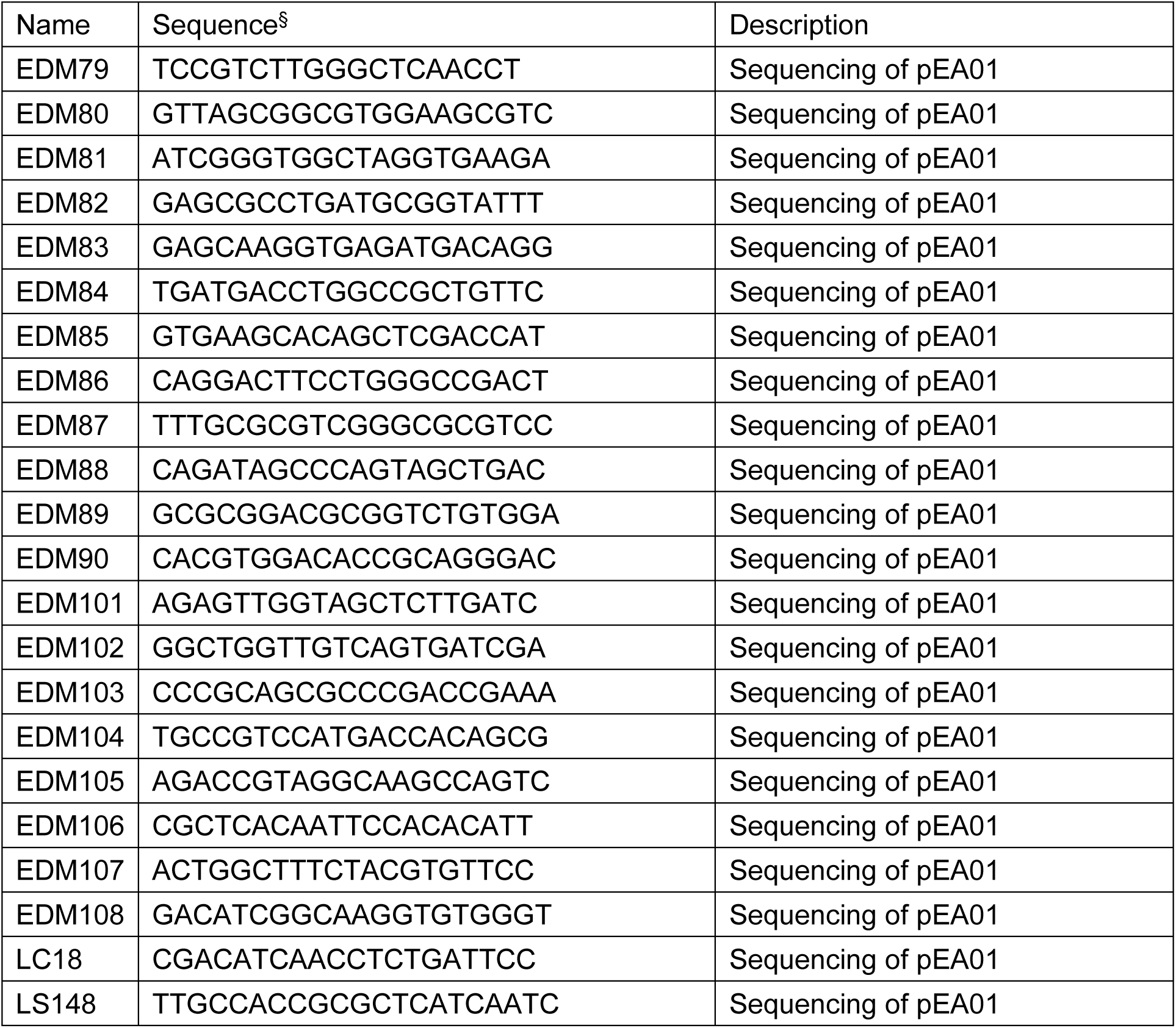
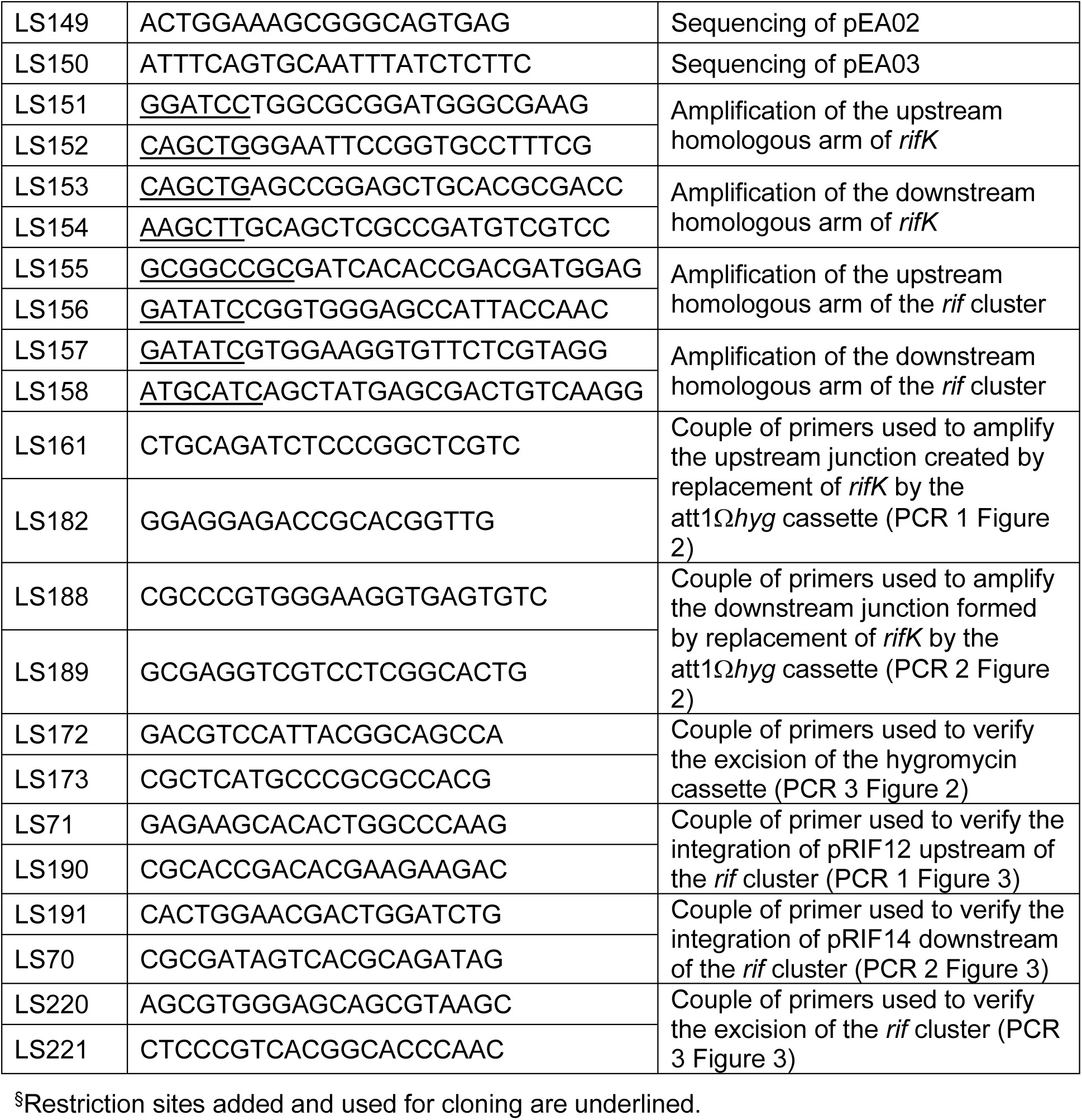
Oligonucleotides used in this study.

## Notes

### Competing Interest Statement

The authors have declared no competing interest.

